# Comparison of picolyl azide-based BONCAT and microautoradiography for assessing the heterotrophic prokaryotic activity in the deep ocean

**DOI:** 10.1101/2025.10.20.683375

**Authors:** Chie Amano, Eva Sintes, Noémie Lebon, Julia Steiger, Danilo Prijovic, Thomas Reinthaler, Ingrid Obernosterer, Kristin Bergauer, Gerhard J. Herndl

## Abstract

Prokaryotes play a central role in marine biogeochemical cycles, yet quantifying their activity requires sensitive methods due to low biomass and metabolic rates, particularly in the deep ocean. One recent method to determine single-cell activity of prokaryotes is bioorthogonal non-canonical amino acid tagging (BONCAT), which offers a non-radioactive approach to measure protein synthesis. However, direct comparisons between BONCAT and radioisotope-based techniques across ocean depth gradients remain limited, particularly for low-activity prokaryotic communities. To address this knowledge gap, we tested an optimised BONCAT protocol using picolyl azide fluorophores (BONCAT-pic) to assess single-cell heterotrophic activity in prokaryotic communities from surface to bathypelagic depths (1000–4000 m) in the Southern Ocean near the Kerguelen Islands. The method was first optimised using aged coastal and open-ocean seawater, and then compared to microautoradiography with ^3^H-methionine uptake. Statistical analysis shows that BONCAT-pic significantly improved detection sensitivity compared to standard azide reagents. BONCAT-pic consistently detected active cells in profiles over the open ocean water column, with cell proportions and fluorescence signals closely correlating with both microautoradiography (R^2^ = 0.9, p < 0.001) and bulk methionine incorporation (R^2^ = 0.6, p < 0.001). Our results demonstrate that BONCAT-pic is a reliable, fluorescence-based method for quantifying heterotrophic activity at the single-cell level, extending its applicability to prokaryotic communities in the deep ocean.

## Introduction

The ocean is a vast reservoir of organic matter [1], and its remineralisation is largely mediated by heterotrophic prokaryotes [2]. These microbial processes are fundamental to oceanic carbon cycling and influence global biogeochemical fluxes [3]. Thus, quantifying the activity of heterotrophic prokaryotes is essential for improving estimates of carbon transformations and turnover across different carbon pools.

One of the most widely used approaches to estimate prokaryotic heterotrophic activity is based on the incorporation of radiolabeled leucine [4–6], a method that provides rapid, sensitive, and specific estimates of prokaryotic carbon biomass production. Leucine uptake measurements are well-suited for routine use due to the manageable incubation volumes and high reproducibility across biological replicates. The technique is also particularly well suited for measurements in bathypelagic waters, where prokaryotic activity and growth rates are substantially lower than those in surface waters [7]. When samples are processed in parallel for microautoradiography, including fluorescence *in situ* hybridisation (FISH), the activity of the prokaryotic community can be visualised and quantified under a microscope by enumerating the uptake of radiolabeled leucine observed as silver grain halos around cells, allowing estimation of group-specific single-cell activity [8].

Methionine is a sulphur-containing amino acid and is essential to all organisms as it universally initiates protein synthesis. While certain taxa, such as the SAR11 clade, are methionine auxotrophs [9], many marine microorganisms are considered capable of *de novo* synthesising methionine [10]. However, since methionine biosynthesis is energetically costly [11], even non-auxotrophs readily assimilate external methionine [12], which makes it useful as a proxy for bulk heterotrophic microbial activity. Uptake of radiolabeled methionine and leucine was compared in the North Atlantic, revealing lower methionine incorporation rates than leucine [13], likely because of the lower proportion of cellular methionine (∼2%) than leucine in marine prokaryotes [4,14].

More recently, bioorthogonal non-canonical amino acid tagging (BONCAT), a non-radioisotope alternative for quantifying protein synthesis, expanded the application of methionine-based approaches to assess microbial carbon production at the single-cell level [15]. BONCAT relies on the incorporation of synthetic methionine analogues, such as L-homopropargylglycine (HPG) or L-azidohomoalanine (AHA), into newly synthesised proteins. These analogues contain alkyne or azide functional groups that can be fluorescently labelled via copper(I)-catalysed azide-alkyne cycloaddition, a reaction commonly referred to as “click chemistry”. The copper catalyst plays a critical role in enabling this reaction, which forms the basis for BONCAT detection via fluorescence microscopy or flow cytometry. Following initial validation on bacterial isolates as well as complex environmental microbial communities [16], BONCAT has been applied to a wide range of aquatic systems and microbes, from viruses [17] to eukaryotic microbes [18], in both pelagic [19] and benthic [20] marine environments. Similar to microautoradiography, BONCAT enables single-cell resolution of metabolic activity, allowing visualisation and quantification of active cells based on fluorescence intensity [19,21]. BONCAT generally correlates well with radiolabeled leucine incorporation [18,19]. However, direct comparisons between BONCAT using HPG and microautoradiography with ^35^S-methionine have shown that the two methods are not fully interchangeable. In particular, BONCAT with the filter-transfer-freeze method to reduce background, exhibits lower sensitivity than microautoradiography, leading to an underestimation of activity [21]. This limitation is especially critical in low-biomass, low-activity samples of the deep ocean, which typically produce weak fluorescence signals.

To reduce the copper-associated toxicity and improve the sensitivity of standard azide tags in BONCAT, picolyl azide fluorophore (BONCAT-pic) was developed. The picolyl group in this fluorophore coordinates copper at the reaction site, accelerating the click reaction and allowing lower, less toxic copper concentrations [22,23]. BONCAT-pic has been applied to microbial communities in hot spring sediments to detect taxon-specific responses to substrate amendments using flow cytometry and cell sorting [24]. Thus, the versatility of general BONCAT combined with the enhanced labelling efficiency of picolyl may offer an effective approach for assessing low-activity prokaryotic communities.

Here, we aimed at evaluating whether BONCAT-pic could serve as a fluorescence-based alternative to microautoradiography for assessing metabolic activity in deep-ocean prokaryotes. We optimised and tested the BONCAT-pic protocol for detecting single-cell protein synthesis activity using aged coastal and open-ocean samples. We then compared the performance to traditional microautoradiography using ^3^H-methionine, based on natural deep-ocean prokaryotic communities collected in the Southern Ocean off the Kerguelen Islands. BONCAT-pic was sufficiently sensitive to detect low-activity cells and yielded results consistent with microautoradiography, demonstrating its reliability as a single-cell approach for assessing heterotrophic activity in deep-ocean prokaryotic communities.

## Materials and Methods

### Sampling and incubation

Seawater samples were collected during several field campaigns in the Atlantic, the Southern Ocean and the Adriatic Sea (see Table 1). To optimize the BONCAT protocol, we used the prokaryotic community of aged seawater (between 3 months and 7 years) collected from 300 m depth in the tropical Atlantic during the GEOTRACES cruise (GEO), from 250 m in the North Atlantic during the MEDEA II cruise (MED), and from 0.5 m depth in the coastal Adriatic Sea off Rovinj, Croatia (ROV). These samples were initially collected with Niskin bottles on board ships (for GEO and MED) or with a bucket (ROV) and stored in acid-washed 25 L polycarbonate carboys.

**Table 1.** Seawater samples used to optimise and test BONCAT. Aged seawater was stored for an extended period at controlled temperature conditions prior to incubation with HPG. Natural prokaryotic community samples were processed immediately after collection on board. IDs are used to indicate the source of samples.

| Sample Type | ID | Location | Cruise /location | Sampling date | Sampling coordinates | Depth (m) | Inc. temp (°C) | Duration (h) | Sample condition |
| --- | --- | --- | --- | --- | --- | --- | --- | --- | --- |
| Aged seawater | GEO | Atlantic Ocean | GEOTRACES (64PE321) | Jul 2010 | Stn 39: 2°N, 41°W | 300 | 4 | 20.4 | Aged for 7 years at inc. temp |
|  | MED | Atlantic Ocean | MEDEA II (64PE356) | Jul 2012 | 50–60°N, 20–40°W | 250 | 4 | 20.1 | Aged for 5 years at inc. temp |
|  | ROV | Coastal Adriatic Sea | Off Rovinj | Nov 2017 | 45.09°N, 13.63°E | 0.5 | 17–20 | 2.1 | Aged for 3 months at inc. temp |
| Natural prokaryotic community | MOB | Southern Ocean | MOBYDICK | Feb–Mar 2018 | Stn M2: 72.02°E, 50.63°S | 50–500 | in situ | 5–15 | Immediately processed |
|  |  |  |  |  | Stn M3: 68.06°E, 50.68°S | 50–1500 | in situ | 6–30 | Immediately processed |
|  |  |  |  |  | Stn M4: 67.21°E, 52.60°S | 100–4000 | in situ | 6–30 | Immediately processed |

Inc. temp: incubation temperature; Duration: duration of incubation with HPG; Stn: Station

The heterotrophic activity of natural prokaryotic communities was assessed in seawater samples from 50–4000 m deep waters in the Southern Ocean during the MOBYDICK cruise. These samples were collected using a conductivity-temperature-depth (CTD) rosette equipped with 12 L Niskin bottles. Contextual physico-chemical data, including microbiology from the MOBYDICK cruise, have been presented elsewhere [25]. Prokaryotic abundance was measured with flow cytometry (FACSCanto II flow cytometer, Becton Dickinson) after fixation with glutaraldehyde (final concentration 1%) and staining with SYBR Green I [26].

Triplicate aliquots, 10–40 mL for each sample, depending on the expected abundance and activity, were distributed into sterile polypropylene centrifuge tubes. One of the aliquots served as a killed control, in which 0.2 µm-filtered formaldehyde (final concentration 2%) was added prior to substrate addition. For incubations, either L-homopropargylglycine (HPG; supplied by the Click-iT HPG Alexa Fluor 488 Protein Synthesis Assay Kit, Thermo Fisher Scientific; final concentration 20 nM) or ^3^H-methionine (L-[methyl-^3^H]-methionine, specific activity: 80 Ci/mmol, ART-0169, Biotrend; final concentration 20 nM) was added to both live samples and killed controls. We also performed parallel incubations with AHA, applying the same concentration and duration; however, the fluorescence signals were substantially lower than those obtained with HPG, and therefore, AHA was not used in subsequent work. The duration of incubations varied with sampling depth: 2–7 h for epipelagic (<200 m), 12– 20 h for mesopelagic (200–1000 m), and 20–30 h for bathypelagic (>1000 m) waters, based on the expected microbial activity at each depth and following the incubation times used in previous leucine-based studies [27]. Incubation times for the aged seawater samples are provided in Table 1. Samples were kept in the dark, either at the storage temperatures of the aged seawater or at in situ temperature ±1°C for samples processed during the cruise. Experiments of HPG and ^3^H-methionine were conducted in parallel for each set of samples using the same volume, incubation times and temperatures. To terminate the incubations, formaldehyde (final concentration 2%) was added to the live samples. Samples were stored at 4°C for 18 h and subsequently filtered onto 0.2 µm white polycarbonate filters (25 mm diameter, Millipore GTTP) using nitrocellulose support filters (25 mm diameter, Millipore HAWP). Filters were rinsed twice with ∼5 mL of Milli-Q water, air-dried, and stored in microcentrifuge vials at –20°C until further processing.

### BONCAT optimization

To optimise the click reaction conditions of the commercially available Click-iT HPG Alexa Fluor Protein Synthesis Assay Kit (Thermo Fisher Scientific), we tested several reagent mixtures. The reaction mix drives the copper-catalysed azide–alkyne cycloaddition to fluorescently label the HPG incorporated by the prokaryotes. Our initial standard protocol followed the click reaction mixture composition in Samo et al. [21] with a copper solution of 2% v/v and an Alexa Fluor 488 azide final concentration of 8 µM. We tested modified versions of this standard reagent mixture (STD), with double the copper concentration (STD 2×Cu) or with double the Alexa Fluor azide concentration (STD 2×AF).

In addition to the standard click reaction reagent, we tested picolyl azide fluorophores from the Click-iT Plus Alexa Fluor 488 Picolyl Azide Toolkit (Thermo Fisher Scientific). Following the manufacturer’s instructions, a copper protectant solution was prepared by mixing copper (II) sulphate (CuSO₄) with the copper protectant at a ratio of 2:1. A CuSO₄/protectant mix was added to the reaction buffer at 2% v/v, followed by Alexa Fluor 488 picolyl azide at 5 µM. This reagent mixture was our standard picolyl azide mixture (Pic). A mixture containing 10 µM of Alexa Fluor 488 picolyl azide was also tested (Pic 2×AF).

For the click reaction, a wedge of one-twelfth of a 25 mm diameter polycarbonate filter was cut with a sterile scalpel and placed into a 1.5 mL microcentrifuge tube containing 300 µL of the reaction mixture. The mix was incubated at room temperature in the dark for 30 min. The filters were washed three times with ∼20 mL of Milli-Q and dried at 37°C for 10 min.

### Cell transfer from filters

Following the click reaction, cells on the filter were transferred to microscope glass slides to reduce background fluorescence and improve the signal-to-background ratio [28]. Several transfer methods were tested: (1) the filter-transfer-freeze technique (FTF), as originally described by Hewes and Holm-Hansen [29] and later used by Samo et al. [21]; we tested both the dry-ice method and cold spray (Cytocool II, 8323, Thermo Scientific); (2) adhesion glass slides (TOMO Adhesion Slides, Matsunami Glass Ind.) and (3) microscope slides coated with a thin layer of adhesive material to promote cell attachment. For this third approach, we tested agarose (2% w/v), gelatine (3% w/v), and polyvinyl acetate glue, a common wood glue, diluted in Milli-Q at a ratio of 1:2 (UHU Holzleim Original, UHU GmbH).

Among these, the gelatine-based method yielded the most effective and consistent cell transfer (see Results section) and was therefore used for subsequent analyses. Briefly, gelatine (Bovine Gelatin, G9391, Sigma-Aldrich) was dissolved in Milli-Q at a final concentration of 3% (w/v) by heating the solution to 43°C for 15 min. Microscope slides were dipped into the warm gelatine. After dipping, one side of each slide was wiped off, and the slides were placed on an ice-cold aluminium plate for 1–5 min to allow the gelatine to solidify. Filter sections were then placed onto the gelatine-coated surface. The slides were left to dry in the dark at room temperature for 20 min to 1 h before gently peeling off the filters. After filter removal, the transferred cells were mounted with a DAPI mixture consisting of 5.5 parts Citifluor (AF1, Citifluor Ltd.), 1 part of Vectashield (Vector Laboratories Inc.), and 0.5 parts phosphate-buffered saline (PBS) containing DAPI at a final concentration of 2 µg mL^−1^. The cells were covered with a cover slip for later counting on the microscope. To monitor potential contamination introduced during the transfer process, a blank membrane filter section was processed in parallel. The detailed protocol is provided in the Supplementary Material.

To normalise fluorescence intensity across samples, InSpeck Green image intensity fluorescence calibration beads (Thermo Fisher Scientific) were initially included as internal standards by applying them directly to the sample after the click reaction (Supplementary material). These beads allow generating a calibration curve over a wide range of fluorescence intensities, from very low (0.3% relative intensity) to the brightest signal (100% relative intensity). However, even the lowest-intensity beads were over-saturated at the exposure time chosen for natural microbial communities, and therefore no calibration curve could be produced. Thus, bead-based normalisation was not applied in this study. Instead, a positive control filter, replicate filters of aged seawater samples with known HPG-positive cell counts and fluorescence intensity was processed alongside the samples as our positive control to ensure internal comparability throughout the analysis. All steps were conducted under dim light to minimise photobleaching of the fluorescent signal. Microscopy images were acquired within several days after mounting. To assess the effect of storage on fluorescent signals after BONCAT, we also evaluated HPG-positive cell counts and fluorescence intensity during storage at –20°C for up to two weeks.

### Microautoradiography

Microautoradiography was performed to assess single-cell methionine uptake, following the protocol described for leucine by Sintes and Herndl [8]. Similar to BONCAT, one-twelfth of each sample filter was used for microautoradiography. Emulsion preparation, exposure, development, and fixing were performed according to the manufacturer’s instructions. The nuclear emulsion (Ilford, Type K5) is light sensitive, thus all steps prior to fixing were carried out in complete darkness. Briefly, filter sections were placed onto glass slides coated with the emulsion. Subsequently, the slides were stored in a light-proof box containing silica gel desiccant and exposed at 4°C for two weeks. Following exposure, slides were developed in developer (Ilford Phenisol Developer) for 4 min, rinsed with Milli-Q, and then fixed (Ilford Hypam Fixer) for 6 min. The filter sections were peeled off, and the cells were mounted with the DAPI mixture described above. Prepared slides were stored at 20°C until counting under a microscope.

### Microscopy and image analysis

BONCAT and microautoradiography slides were examined using an epifluorescence microscope (Axio Imager M2, Carl Zeiss) equipped with Zeiss filter sets for DAPI (Filter Set 49, excitation G 365, emission BP 445/50) and Alexa Fluor 488 (Filter Set 44, excitation BP 475/40, emission: BP 530/50), as well as transmitted light. Images were acquired at 1000-fold magnification using a CCD-based monochrome digital camera (AxioCam MRm, Carl Zeiss). At least 10 fields of view were recorded per filter section, where each pixel on the image corresponded to 0.01 µm^2^. The exposure time of DAPI and transmitted light images was auto-adjusted. Images of the Alexa Fluor 488 filter channel were captured with a fixed exposure time of 2000 ms.

HPG-positive cells were analysed using the Automated Cell Measuring and Enumeration tool (ACMEtool 3.0, Bennke et al [28]). We aimed at counting a minimum of 800 DAPI-stained cells. The accuracy of cell recognition by ACMEtool and cell counting software is influenced by the signal-to-background ratio. Although signal detection improved after the cell transfer step, all images were manually inspected to confirm proper cell detection. Cells with Alexa Fluor 488 and DAPI signals were considered HPG-positive. Killed controls were used to define detection thresholds and to set the baseline for Alexa Fluor 488 signal intensity, corresponding to <2% HPG-positive cells among the total DAPI-stained prokaryotes, following the threshold used in a previous study [19]. This low background observed in the killed controls also indicated negligible interference from phytoplankton autofluorescence being in a similar wavelength as the Alexa Fluor dye. Single-cell fluorescence from the HPG was calculated as the mean grey value (MGV) multiplied by the area of the HPG-positive signal (µm²), divided by the incubation time with HPG (T, in hours). The MGV is defined as average pixel brightness within the detected cell area on an 8-bit scale, where 0 = black and 255 = white. Finally, the total HPG-derived fluorescence was calculated by summing the cell-specific values and expressed in MGV µm^2^ L^−1^ h^−1^.

For microautoradiography, the silver grain halo surrounding DAPI-stained cells was analysed using the AxioVision SE64 imaging software (Rel 4.9, Carl Zeiss) with a custom-made macro to measure the halo size associated with DAPI-stained cells. Similar to BONCAT, the single-cell radiolabeled methionine incorporation was estimated based on the silver grain halo area, expressed as µm^2^ cell^−1^ h^−1^. The total incorporation was calculated by summing all halo areas, expressed as µm^2^ L^−1^ h^−1^.

### Bulk methionine and leucine incorporation

Bulk substrate incorporation rates were determined using the filtration method [30]. Incubation and fixation of methionine incubations followed the procedure as described above using one killed control and two live samples per depth. For leucine incorporation, incubations were conducted with [3,4,5-^3^H]-L-leucine (specific activity: 120 Ci mmol^−1^, ART-0470, Biotrend) at a final concentration of 10 nM (≤200 m) and 5 nM (>200 m). Three live samples and two formaldehyde-killed controls (2% final conc.) were prepared per depth. Following incubations at in situ temperature in the dark, live samples were fixed with formaldehyde and filtered through 0.2 µm polycarbonate filters (25 mm diameter, Millipore GTTP) using nitrocellulose support filters (Millipore HAWP). The filters were overlaid and rinsed twice with 5% ice-cold trichloroacetic acid for 5 min to precipitate proteins and subsequently rinsed once with Milli-Q. The filters were dried, transferred to 20 mL scintillation vials and 8 mL of scintillation cocktail (FilterCount, Perkin Elmer) was added. Disintegrations per minute (DPM) were measured using a calibrated liquid scintillation counter (Tri-Carb 2100, Perkin Elmer) and converted to methionine or leucine incorporation rates, expressed as pmol methionine or leucine L^−1^ h^−1^.

### Analysis and presentation

All statistical analyses and data visualisation were performed in R version 4.4.2 [31] using the following R packages: tidyverse [32], ggsignif, rstatix, readxl, data.table, viridis, marmap, and ggspatial.

## Results

### Evaluation of the cell-transfer method and sample handling

Several modifications were introduced to the previous click chemistry protocol [21], improving reaction efficiency and enhancing fluorescence signal detection associated with HPG incorporation. To reduce background fluorescence caused by non-specific deposition of fluorophores on the membrane filter, we evaluated methods for transferring cells from filters to glass slides using aged seawater samples (Fig. 1A).

**Fig. 1.**
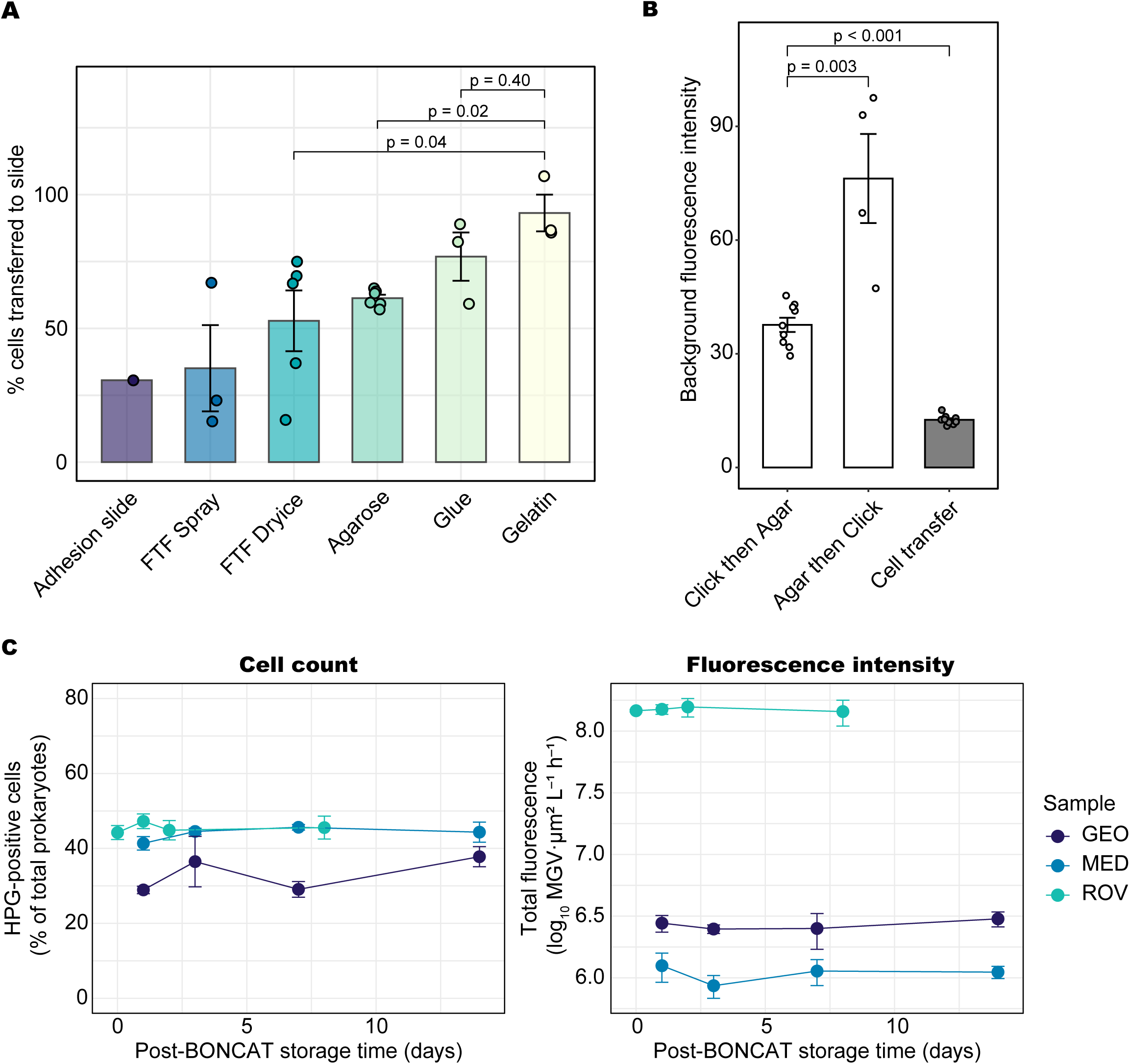
Effects of cell transfer and storage conditions on BONCAT sample integrity. (A) Comparison of cell-transfer methods using aged seawater samples (ROV), where filters were embedded in agarose without BONCAT. (B) Background fluorescence intensity measured after BONCAT in aged seawater samples (GEO and MED). ‘Agar then Click’ indicates filter samples embedded in agarose before the click reaction; otherwise, the click reaction was performed first. ‘Cell transfer’ indicates the background intensity of slides where cells were transferred. (C) Relative percentages of HPG-positive cell counts and their fluorescence intensities in aged samples (ROV, MED, GEO) after BONCAT and storage at –20°C. Statistical differences were evaluated with the Wilcoxon rank-sum test; p-values are indicated in panels A and B.

We tested the cell transfer methods (FTF, adhesion glass slides, coating with adhesive material. We used filter sections already embedded in 0.1% low-melting-point agarose (A9414, Sigma-Aldrich), considering the potential integration with catalysed reporter deposition fluorescence in situ hybridisation (CARD-FISH) described in Teira et al. [33]. We tested whether cells can still be efficiently transferred to glass slides, despite the presence of the agarose layer. Our results showed that, although the FTF method and adhesion slides are clean and simple procedures, both methods resulted in low cell recovery after filter transfer (adhesion slide: 31% recovery, n = 1; FTF with cold spray: 35 ± 28%, n = 3; FTF with dry ice: 53 ± 25%, n = 5; mean ± SD, Fig. 1A). The highest recovery was achieved using gelatine (93 ± 12%, n = 3) and glue (77 ± 16%, n = 3). There was no statistically significant difference between these two methods (Wilcoxon rank sum test, p = 0.4, n = 6). Neither gelatine nor glue interfered with DAPI staining, and blank filters processed alongside the samples showed no signs of contamination from the coating solutions checked under the microscope. However, we selected gelatine for further use due to its easier handling than glue and resulting in a more uniform focal plane of the slide, which ensures high quality images.

To assess background fluorescence unrelated to the cellular signal, we analysed filters after the click reaction with a standard azide fluorophore. When cell transfer using gelatine was performed, background fluorescence was significantly reduced by approximately threefold (Wilcoxon rank sum test, p < 0.001, n = 18) and the MGV decreased from 38 ± 6 to 13 ± 1 (n = 9), indicating that a substantial portion of the observed background originated from the membrane filter and fluorophore residues, with the apparent intensity likely enhanced by light scattering from the underlaying white polycarbonate surface (Fig. 1B). High background fluorescence can mask low levels of HPG incorporation, thus reducing background is important for detecting weak BONCAT signals. We noted that on filters already embedded in agarose prior to performing the click reaction, the MGV was increased to 76 ± 24 (n = 4), indicating substantial background fluorescence (Wilcoxon rank sum test, p = 0.003, n = 13, Fig. 1B). These results confirm previous recommendations to perform the click reaction prior to FISH [34].

Through the cell transfer, a heterogeneous mixture of cells in terms of size, structure, and morphology has to be transferred to the glass slides. Interestingly, the combined count of transferred and remaining cells occasionally exceeded the initial abundance by 10–20%, regardless of the transfer method used, likely due to overcounting caused by partial cell fragmentation during filter peeling. Nevertheless, when gelatine was used, more than 90% of cells remained morphologically intact, indicating that fragmentation and cell loss had minimal impact on activity estimates.

We also tested the effect of sample storage after BONCAT to evaluate whether immediate microscopic analysis is necessary. Multiple identical filters were prepared in parallel using aged samples and processed through the click-reaction step. The filters were mounted on four separate slides and examined after different storage durations (at T1–T4), to avoid repeated light exposure during microscopy. Images were taken for all slides under identical settings and analysed. Our results showed that storage at –20°C for up to two weeks did not affect HPG-positive cell counts or fluorescence intensity (Friedman rank sum test, p > 0.05 for both cell-count and fluorescence intensity; Fig. 1C).

### BONCAT protocol optimisation: Alexa Fluor azide vs. picolyl azide

We tested several BONCAT click-reaction mixtures using either standard Alexa Fluor azide (STD) or picolyl azide (Pic), with varying concentrations of copper and fluorophore (see Methods). The comparison was conducted using aged seawater from the GEO and natural microbial communities collected at 100 m depth from MOBYDICK (stations M2 and M4; Table 1). We compared the percentage of HPG-positive cells among total DAPI-stained cells and the total HPG-derived fluorescence signal. All comparisons were made relative to the standard azide-based protocol (STD) to evaluate the differences between the protocols.

No significant differences were observed among standard azide conditions (STD, STD 2×Cu, STD 2×AF) for HPG-positive cell percentages or total fluorescence intensity (Kruskal–Wallis test, p > 0.05, n = 9), whereas the use of the picolyl azide significantly increased both metrics compared to the standard reaction mix (STD vs. Pic and STD vs. Pic 2×AF; Wilcoxon rank sum test, one-sided, p = 0.03 for both cell counts and intensity, n = 6). Micrographs showed lower background intensity in Pic compared to STD (Fig. 2C), improving contrast between signal and background with picolyl azide and detection of weak fluorescence. Varying Alexa Fluor azide concentration had little effect on the HPG-positive cell percentages but slightly increased intensity in the Pic 2×AF condition (Fig. 2B).

**Fig. 2.**
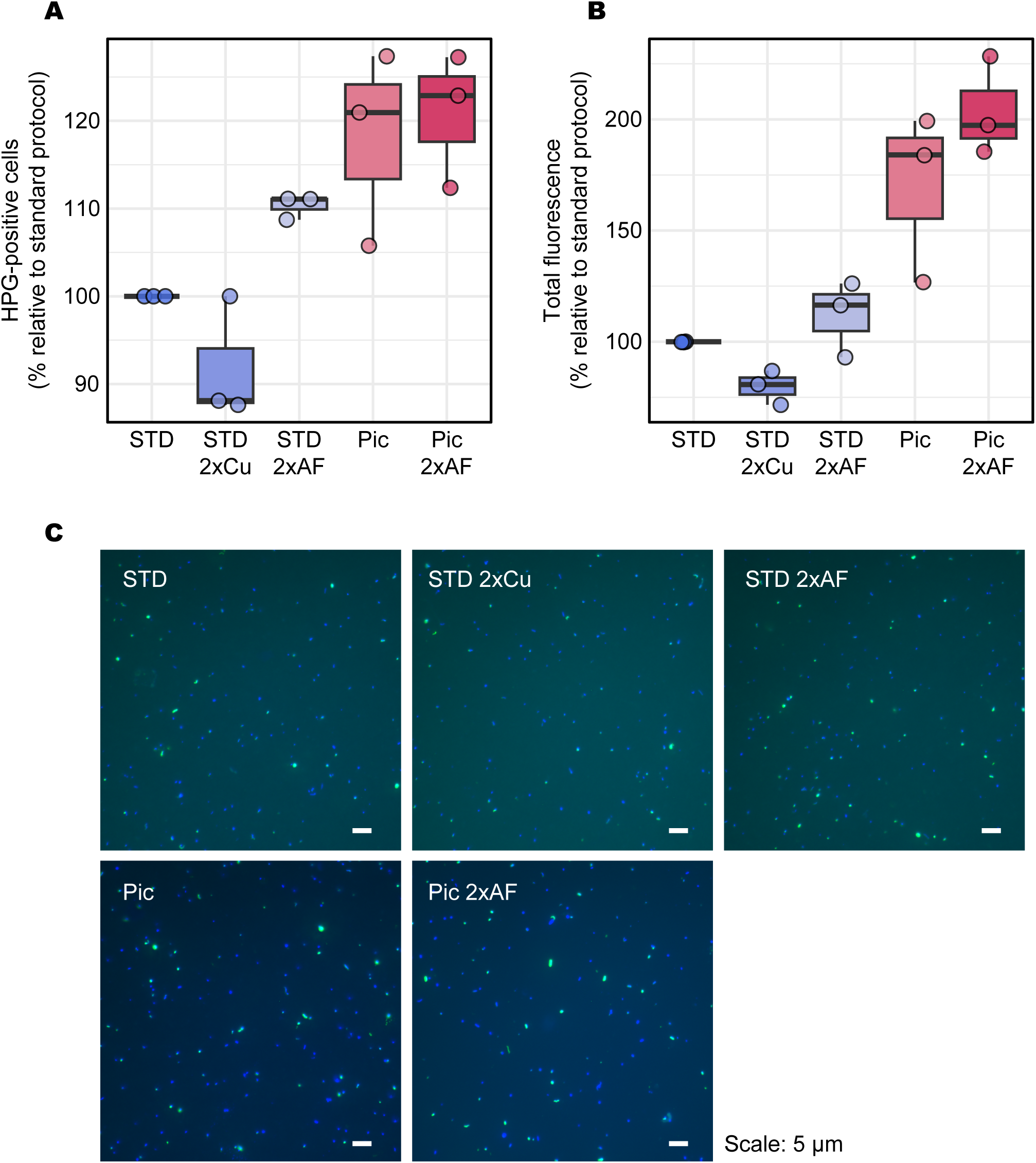
Optimisation of the click reaction using the standard and picolyl azide protocol. Comparison of protocols of the standard azide fluorophore (STD) and picolyl azide fluorophore (Pic). (A) Percentage of HPG-positive cells and (B) Fluorescence intensity of HPG-positive cells expressed relative to the protocol STD. (C) Epifluorescence micrographs of natural microbial communities incorporating HPG at 100 m depth off Kerguelen (MOB, Stn M4).

Taken together, these results suggest that our BONCAT protocol using picolyl azide offers enhanced sensitivity and improved signal-to-background contrast compared to the previous standard azide-based approach. Based on these findings, we selected Pic 2×AF as the optimised protocol for subsequent comparisons with the standard azide (hereafter referred to as BONCAT-pic and BONCAT-std, respectively). The detailed protocol of BONCAT-pic is provided in the Supplementary Material.

### Bulk prokaryotic abundance and substrate incorporation off the Kerguelen Islands

To evaluate the applicability of BONCAT for determining protein synthesis activity in natural prokaryotic communities, we compared the BONCAT-std and BONCAT-pic on samples collected at three stations (M2, M3 and M4) with depths ranging from 50 m to 4000 m during an expedition in the Southern Ocean off Kerguelen Islands (Fig. 3A, Table 1). Station M2 was located on the plateau (bottom depth: 520 m), while M3 and M4 were located off the plateau, with bottom depths of 1730 m and 4370 m, respectively.

**Fig. 3.**
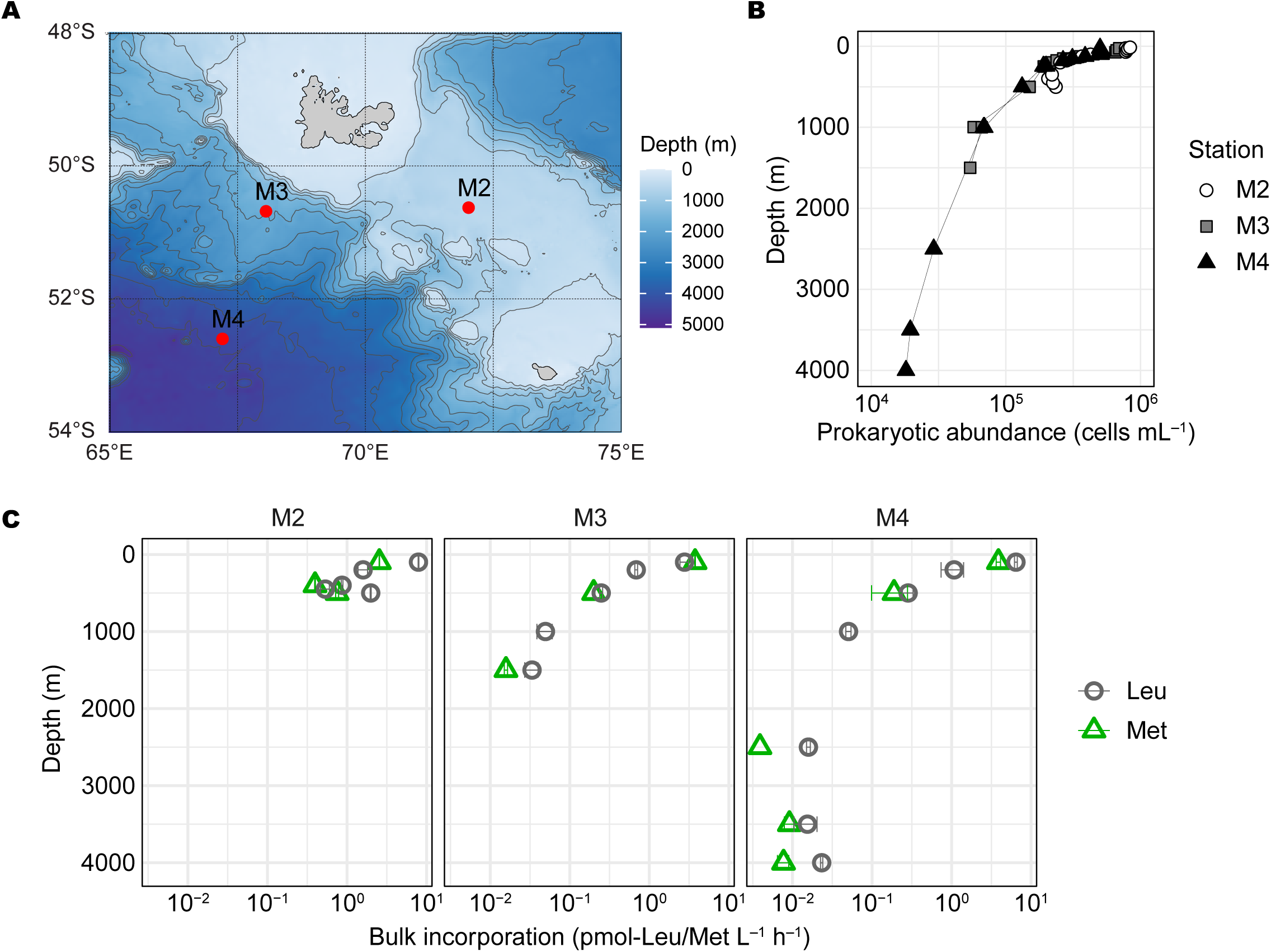
Vertical profiles of prokaryotic abundance and heterotrophic prokaryotic activity in the Southern Ocean. (A) Station map around the Kerguelen Islands. Sampling stations are indicated as red circles labelled with station names; background contours show bathymetry. (B) Depth profiles of prokaryotic abundance and (C) methionine and leucine incorporation rates at three stations. Error bars represent standard deviations.

Prokaryotic abundance showed a typical open ocean profile, decreasing exponentially from 5–8 × 10^5^ cells mL^−1^ at 25 m depth to about 2 × 10^4^ cells mL^−1^ at ∼4000 m depth (Fig. 3B). Similarly, bulk incorporation rates of radiolabeled methionine and leucine decreased with depth, reflecting the decline in prokaryotic abundance (Fig. 3C). Bulk ^3^H-methionine incorporation rates were highest in the epipelagic (3.4 ± 0.7 pmol L^-1^ h^-1^, n = 3) and 0.01 ± 0.005 pmol L^-1^ h^-1^ (n = 4) in the bathypelagic waters. Comparing the methionine and leucine incorporation rates, leucine uptake was consistently higher than methionine uptake (paired Wilcoxon signed-rank test, p = 0.02, n = 11), in agreement with a previous study from the Atlantic Ocean [13]. Additionally, at stations M3 and M4, the difference between methionine and leucine incorporation rates became more pronounced with depth (Spearman rank correlation, ρ = 0.77, p = 0.025, n = 8).

### Single-cell analysis of prokaryotic activity using BONCAT and microautoradiography

To assess prokaryotic activity at the single-cell level, we conducted analyses using both BONCAT and microautoradiography. For the comparison between BONCAT-std and BONCAT-pic, samples of stations M2 and M4 were analysed. When applying BONCAT-std, the percentage of HPG-positive cells remained relatively constant throughout the water column at station M2 (27 ± 3%, n = 6), while at station M4, it decreased from ∼30% in the epipelagic layer to 8 ± 6% (n = 4) in the bathypelagic (Fig. 4A). When BONCAT-pic was used, detection rates nearly doubled across depths, resulting in 52 ± 4% HPG-positive cells at station M2 (n = 7). At stations M3 and M4, the HPG-positive percentages using BONCAT-pic were 52 ± 2% (n = 3) in the epipelagic, 32 ± 5% (n = 3) in the mesopelagic, and 18 ± 11% (n = 5) in the bathypelagic layers (Fig. 4A). These values from BONCAT-pic were essentially the same as the percentages of methionine-positive cells determined via microautoradiography (Fig. 4A). At depths >2000 m, however, the relative abundance of HPG-positive cells detected by BONCAT-pic tended to be lower than that detected by microautoradiography (Fig. 4A and B). Additionally, the variation between technical replicates was slightly higher in BONCAT than in microautoradiography (Fig. 4B). Overall, BONCAT-pic exhibiting an approximately 1:1 relationship with a root mean square deviation of 3.5%, compared to 15% for BONCAT-std (Fig. 4B).

**Fig. 4.**
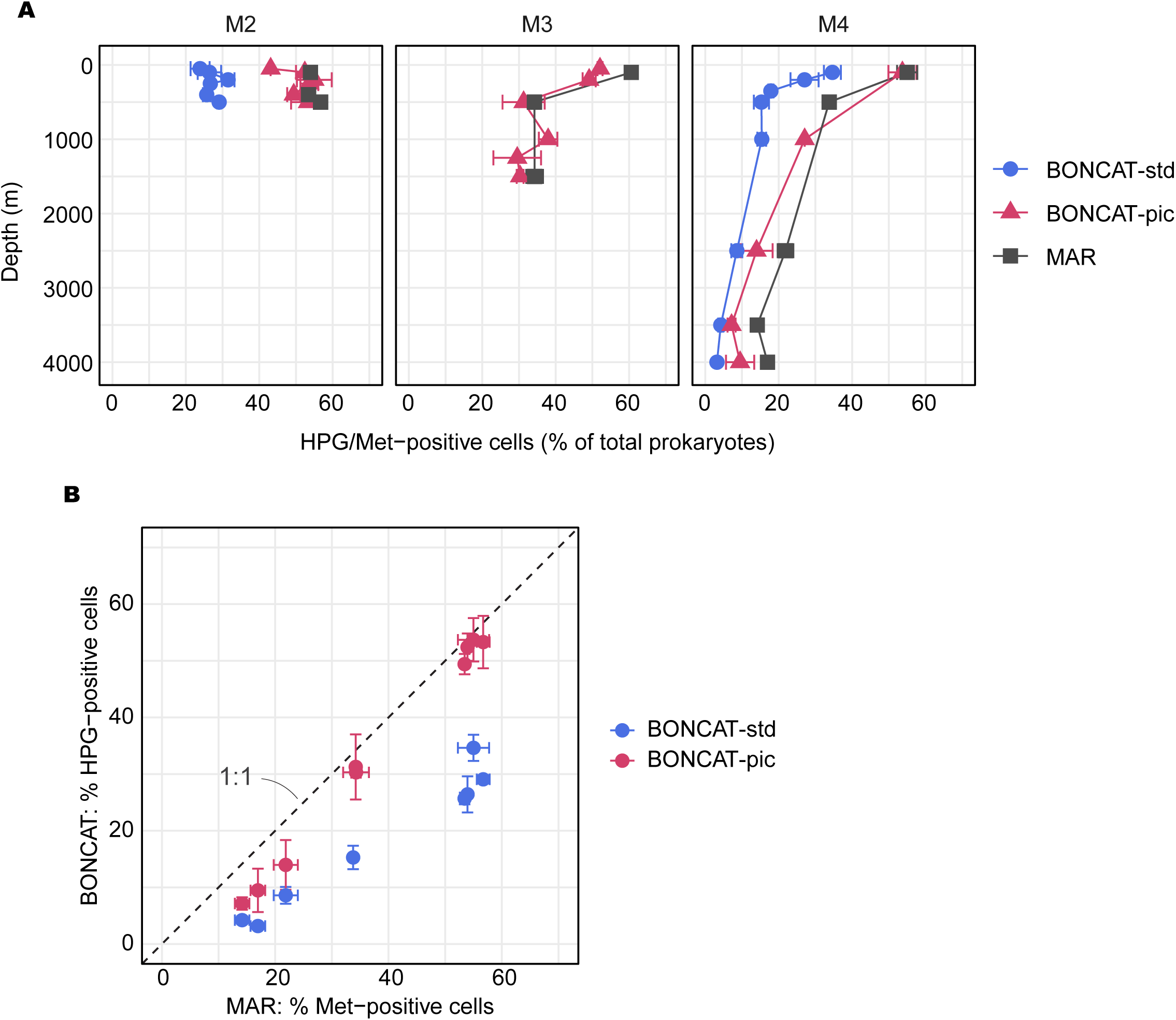
Comparison of standard azide (BONCAT-std) and picolyl azide (BONCAT-pic) fluorophores with microautoradiography (MAR). Samples were collected from the epipelagic to bathypelagic waters during the MOBYDICK cruise and incubated with either HPG or ³H-methionine under identical concentrations, temperatures, and incubation times. (A) Vertical profiles of HPG or ^3^H-methionine positive cells detected by BONCAT and microautoradiography (MAR). (B) Comparison of the percentage of HPG and ^3^H-methionine-positive cells between MAR and BONCAT protocols across depths.

To detect weak fluorescence signals, a fixed long exposure time (2000 ms) was used across all samples. This improved detection in deep-water samples but caused signal saturation, MGV of close to 255, in a few of highly active cells, mostly from the epipelagic layer. However, only 0.4% of HPG-positive cells showed MGV >225, indicating minimal signal saturation. Some cells contained more than one HPG signal. However, such overestimation was rare, affecting only 1 ± 3% of signals (n = 32).

### BONCAT-pic fluorescence intensity in open ocean samples

In addition to cell count–based comparisons, we analysed BONCAT-pic fluorescence signal intensity at both the community and single-cell level. These measurements were compared to the total silver grain halo area obtained by microautoradiography as well as to the bulk ^3^H-methionine incorporation rates.

Total HPG-derived fluorescence intensity showed a strong positive correlation with both total silver grain halo area (R^2^ = 0.86, p < 0.001; Fig. 5A) and bulk ^3^H-methionine incorporation (R^2^ = 0.61, p < 0.001; Fig. 5B). Single-cell activity decreased with depth in both HPG and methionine uptake (Fig. 5C). At station M4, cell-specific activity decreased by nearly one order of magnitude from the epipelagic to bathypelagic layer, consistent with the depth-dependent decrease observed in bulk prokaryotic activity (Fig. 3C). The relative frequency distribution of single-cell HPG fluorescence intensity obtained with BONCAT-pic closely matched that of ^3^H-methionine uptake measured by microautoradiography, with comparable single-cell activity range of approximately two orders of magnitude.

**Fig. 5.**
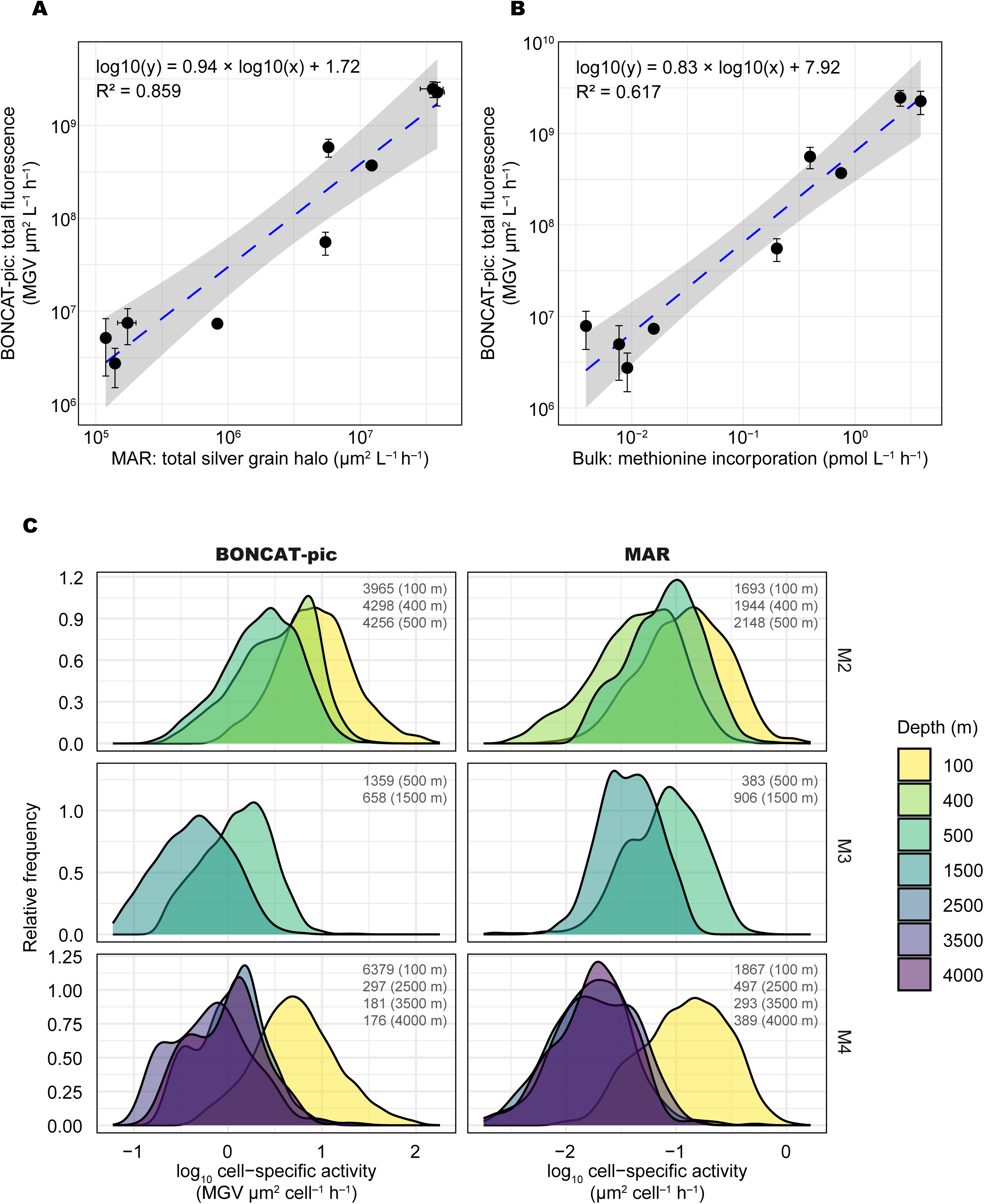
Fluorescence-based analysis of prokaryotic activity using MOBYDICK samples. (A) Total fluorescence from BONCAT-pic versus total silver grain halo area from microautoradiography (MAR) and (B) versus bulk methionine incorporation rates. Dashed blue lines indicate regressions on log_10_ transformed data; gray shading mark 95% confidence intervals. (C) Distribution of single-cell substrate uptake activity, shown as probability density plots based on BONCAT-pic and MAR. These plots represent the relative frequency distribution of active cells across a range of cell-specific substrate uptake rates. The number of cells analysed and the corresponding sampling depths are indicated in each panel.

## Discussion

Although picolyl azide reagent has been commercially available for over a decade, the BONCAT-pic has not been quantitatively evaluated for environmental microbial communities, particularly in the deep ocean. This is mainly because direct validation requires parallel comparisons with radioisotope-based assays, which are technically demanding and often subject to regulatory restrictions. There is no leucine analogue compatible with click chemistry, restricting BONCAT to methionine-based labelling. Improving the sensitivity of detecting the methionine uptake at the single-cell level is essential for quantifying microbial activity in the deep ocean, where metabolic rates are low. In this study, BONCAT-pic significantly increased the detection of HPG-positive cells compared to BONCAT-std and produced results comparable to microautoradiography and bulk methionine incorporation. These findings demonstrate that the enhanced sensitivity of BONCAT-pic is well-suited for assessing single-cell methionine uptake in open-ocean samples. Because BONCAT-pic requires only standard laboratory infrastructure, its accessibility makes it a practical and widely applicable approach for studying microbial communities with low metabolic activity.

Our results show that the choice of azide significantly affects the detection of HPG-positive cells. Picolyl azide consistently revealed a higher proportion of active heterotrophic cells across all depths compared to the standard azide (Fig. 4). Because HPG incubation and fixation conditions were identical, this difference likely originates from the click reaction. The copper-chelation properties of picolyl enhance the efficiency of the copper-catalysed azide–alkyne cycloaddition [22], increasing labelling efficiency and fluorescence intensity. Consequently, the proportion of HPG-positive cells was approximately doubled throughout the water column when using picolyl azide.

We used a final concentration of 20 nM HPG in all water column incubations. This is higher than typical concentrations of dissolved free methionine in the oligotrophic ocean, ranging from <1 nM to ∼3 nM [35,36]; but is lower than the concentrations used in previous BONCAT studies (∼1 µM) [19]. Generally, elevated substrate concentrations have two opposing effects: they can suppress biosynthesis of the target compound, reducing dilution of the labelled pool, while simultaneously altering substrate uptake rates in prokaryotes [30,37]. Therefore, the measured signal should be interpreted as potential uptake rates rather than in situ activity. Because amino acid uptake is concentration-dependent [37,38], we adjusted the HPG concentration to minimise artificially stimulating uptake while ensuring detectable labelling. Using similar nanomolar concentrations for HPG, methionine and leucine also facilitated comparisons between methods.

A key assumption of BONCAT is that incorporation of the synthetic methionine analogue HPG does not affect cellular metabolism. Although HPG is generally considered non-toxic, previous studies have reported concentration-dependent effects: high HPG concentrations (in the µM level) inhibited growth in *Escherichia coli*, unlike ^3^H-methionine [39], and induced stress response in cyanobacteria [40]. Given the physiological diversity of marine prokaryotes, predicting such toxic effects in natural assemblages is difficult. Nevertheless, the BONCAT-pic protocol employed here used HPG at nM concentrations, which are well below the toxicity thresholds reported in laboratory studies. Moreover, HPG uptake patterns were comparable to those of ^3^H-methionine in deep-ocean microbial communities (Fig. 4B), suggesting no apparent limitation and inhibition in synthetic substrate uptake. This also likely reflects the low ambient methionine concentrations in the deep ocean, where cells readily incorporate available analogues such as HPG.

Microautoradiography remains the standard for assessing single-cell metabolic activity, despite increasing regulatory and logistical constraints associated with radioisotope use. In this study, microautoradiography consistently detected the highest percentage of active heterotrophic cells across all stations, indicating that BONCAT, particularly with the standard azide fluorophore, may underestimate active cell populations in oligotrophic environments. In contrast, BONCAT-pic showed strong correlation with microautoradiography, although detection rates were somewhat reduced in lower bathypelagic samples (> 2000 m). Because only a small number of lower bathypelagic samples (n = 3) was analyzed, this trend should be interpreted with caution. However, this apparent difference could be either due to weak fluorescence signals near the detection threshold or actual differences in uptake between methionine and HPG.

Converting fluorescence intensity to absolute substrate uptake rates requires careful consideration, as measurements can be influenced by factors such as signal saturation, background fluorescence, substrate concentration and incubation time. Selecting an appropriate exposure time for microscopy effectively minimises these artefacts. Similar to the previous work [41], an additional advantage of BONCAT-pic combined with epifluorescence microscopy is its ability to localise fluorescence within individual cells, allowing visualisation of active protein synthesis in relation to cell morphology and spatial organisation (e.g., particle-attached or dividing cells). Moreover, taxon-specific growth rates in natural communities can be estimated from the frequency of dividing cells, as demonstrated by Brüwer et al. [42]. Integrating BONCAT-pic with such morphological or taxon-specific analyses will advance our understanding of heterotrophic activity of deep-sea microbes.

## Conclusion

This study demonstrates that BONCAT with picolyl azide results in a similar number of metabolically active cells as microautoradiography across a broad range of oceanic depths. The improved detection efficiency of picolyl azide likely resulted from its copper-chelating properties, which enhance the click-reaction efficiency and fluorescence signal strength, particularly for low-activity cells. Thus, BONCAT-pic represents a robust method for quantifying methionine-based protein synthesis at the single-cell level in natural microbial assemblages, facilitating a deeper mechanistic understanding of microbial processes in the pelagic ocean.

## Acknowledgements

We thank the captains and crews of the MOBYDICK, MEDEA II, and GEOTRACES cruises aboard R/V *Marion Dufresne* and R/V *Pelagia* for their assistance at sea. We also thank F. Malfatti for the lab workshop introducing the BONCAT protocol and P. Catala for flow cytometry analyses.

## Conflicts of interest

The authors declare no conflict of interest.

## Funding

This work was supported by the Austrian Science Fund (FWF; projects I486-B09, Z 194-B17, P35587-B) and the European Research Council (ERC) Advanced Grant (projects MEDEA 268595 and NEREIDES 101141581) to G.J.H. C.A. received funding through a Helmholtz Visiting Researcher Grant from the Helmholtz Information & Data Science Academy (HIDA). E.S. was supported by the Spanish Research Agency (PID2020-118877GB-I00/ MICIU/AEI/473 10.13039/501100011033). T.R. was supported by the Austrian Science Fund (FWF; project BASS I5942-N, PADOM P23221-B11). The MOBYDICK– THEMISTO cruise (https://doi.org/10.17600/18000403) was supported by the French oceanographic fleet (“Flotte océanographique française”), the French ANR (“Agence Nationale de la Recherche”, AAPG 2017 program, MOBYDICK Project number: ANR-17-CE01-0013), and the French Research program of INSU-CNRS LEFE/CYBER (“Les enveloppes fluides et l’environnement” – “Cycles biogéochimiques, environnement et ressources”).

## Data availability

Metadata from the MOBYDICK cruise can be found on https://doi.org/10.17600/18000403. Detailed protocols for the BONCAT-pic are provided in the Supplementary Material. Other data can be provided upon request to the corresponding author.

## Notes

### Competing Interest Statement

The authors have declared no competing interest.

