## Supplementary Material for "Comparison of picolyl azide-based BONCAT and microautoradiography for assessing the heterotrophic prokaryotic activity in the deep ocean"

### BONCAT-pic protocol for pelagic prokaryotes (for microscopy)

### Sampling, incubation, and fixation

1. Check the cell abundance in samples to estimate the appropriate volume (Table 1).
2. Transfer seawater samples to 50 ml centrifuge conical tubes (two tubes: one for the live sample and one for the killed control). Fix the control tubes with 0.2 µm-filtered formaldehyde (final concentration 2%) and wait for 15 minutes.
3. Add substrate for both samples and killed controls, for example: 20 nM final concentration of HPG for open ocean samples. However, note that the concentrations are highly dependent on the sample; a prior check with concentration kinetics is recommended.
4. Incubate in the dark at in situ temperature. Take into account the expected activity of your sample (for example, incubation times for pelagic prokaryotes with HPG are: epipelagic: 2-7 h, mesopelagic: 12-20 h, bathypelagic: 20-30h, which are slightly longer than the leucine-based assay; time-course kinetics are preferred before starting the experiment).
5. Terminate the live samples with 0.2 µm-filtered formaldehyde (final conc. 2%) and store at 4°C in the dark for 1 to a maximum of 24 h.
6. Filter the sample onto white polycarbonate filters (Millipore, GTTP, 25mm diameter), using a nitrocellulose support filter (Millipore, HAWP).
7. After filtering the sample, wash the filter twice with ~5 mL of MilliQ water.
8. Air-dry filters. Store at -20°C until processing. Filters can be kept frozen for several months.

**Table 1. Example of filtration volume**

| Cell abundance in seawater | Volume to filter (ml) |
|---|---|
| $2x10^4$ cell $ml^{-1}$ | 100 |
| $5x10^5$ cell $ml^{-1}$ | 10 |
| $1x10^6$ cell $ml^{-1}$ | 5 |

### Click-chemistry

**based on Samo et al. (2014), adapted for picolyl azide:**

*Work under dim light conditions

1. Thaw Sigma water, buffer additive, and Alexa 488 picolyl azide at room temperature in the dark. Keep the buffer additive on ice once it's thawed.
2. Cut filter sections into 1/12 parts of a 25mm diameter filter and write a label on each filter section with a pencil. Additionally, cut a filter sample with a known HPG-positive cell count as a positive control, and also cut a blank new filter as a negative control.
3. Prepare the reaction buffer (see Table 2).
4. Incubate filter sections in the reaction buffer at RT in the dark for 30 min.
5. After incubation, wash 3x in excess Milli Q water.
6. Place the filter sections on blotting paper (with the sample side up) and dry them at 37°C in a hybridization oven for 10 min. After the click reaction, avoid exposing the sample directly to intense light; you may work in a room with dim illumination.

**Table 2. Reaction buffer mixture**

| Stock reagent | Vol (µl)* | Vol (µl)** |
|---|---|---|
| Sigma water | 154 | 231 |
| 10x reaction buffer | 20 | 30 |
| Copper protectant | 4 | 6 |
| Alexa 488 picolyl azide | 2 | 3 |
| 10x buffer additive | 20 | 30 |

* total 200 µl in 0.6 ml tube (~10 filter sections)
**total 300 µL in 1.5 ml tube (~15 filter sections)

### FISH or CARD-FISH

If combining BONCAT with FISH or CARD-FISH, perform FISH or CARD-FISH protocol at this stage. Begin from the embedding step. Note that embedding should not be performed before the click reaction; otherwise, you will encounter very high background noise.

### Internal standard

* This is optional. For low-activity cells, such as pelagic prokaryotes, the bead calibration curve will far surpass their natural community signal intensity; therefore, this step can be omitted.

1. Sonicate the bead solution (0.3% intensity, 1:10 dilution with Sigma water) for 10 min.
2. Immediately after sonication, pipette 5 µl drops (one drop for each filter section) of the bead solution onto clean cover slips (10 drops per coverslip) and place the filter sections on top of the bead drops (with the sample side down).
3. Let the filters dry at 37°C in a hybridization oven for 15 min in the dark.
4. After drying, carefully remove the filters from the cover slip.

### Cell transfer

1. Prepare the gelatin solution in a 50 mL conical centrifuge tube (see Table 3).
2. Warm the gelatin solution to 43°C in the water bath for 15 min until it dissolves.
3. Dip a slide glass into the gelatin solution to coat it.
4. Wipe off the gelatin only from the back side of the slide. Place the gelatin-coated slide on an ice-cold aluminium plate for 1-5 min to solidify. The time depends on the laboratory's humidity. Check carefully to make sure it doesn't dry out.
5. Place the sample filter sections on the gelatin-coated slide with the sample side facing down (max 12 pieces/slide). Add a new filter section as a control for the gelatin solution.
6. Dry the slide at RT for 20min–1h in the dark, until the gelatin is completely dry.
7. Mark the filter's location with a permanent marker. Write down the sample ID.
8. Gently wet the edge of the filter with an MQ-wetted cotton swab, if necessary. Carefully hold the wet edge with forceps and gently peel the filter away from the slide.
9. Mount with DAPI mix and store at -20°C until taking images with a microscope. Typically, images are captured within 24 h of completing the filter section processing.

**Table 3. Gelatin solution**

| Reagent | Volume | Final conc. |
|---|---|---|
| Gelatin | 0.6 g | 3% |
| MQ | 20 mL | |

### 2. Buffers and Chemicals

**10x Buffer additive**

1. Add 2 mL of Sigma water to the bottle (Component E) and mix until completely dissolved.
2. Make 100-200 µL aliquots in sterile 0.6 mL microcentrifuge tubes.
3. Store the aliquots at ≤–20°C. This solution is stable for up to 1 year.

| Stock reagent | Volume | Final conc. |
|---|---|---|
| Buffer additive (Component E)<br>Sigma water | powder<br>2 ml | 10x |

**Alexa Fluor picolyl azide (PCA) stock (1000 µM)**

Make ~50µL aliquots (see Table below) in black sterile microcentrifuge tubes and store at -20°C.

| Stock reagent | Volume | Final conc. |
|---|---|---|
| Alexa 488 PCA (Component A)<br>DMSO | 1 tube<br>105 µL | 1000 µM |

**CuSO4 copper protectant pre-mix**

| Stock reagent | Volume | |
|---|---|---|
| $CuSO_4$ (Component C)<br>Copper protectant (Component D) | 200 µL<br>100 µL | 2:1 |

**DAPI mix 2 µg/ml**

| Stock reagent | Volume (µl) | Final conc. |
|---|---|---|
| DAPI 50µg/ml | 40 | 2 µg/ml |
| 1xPBS | 70 | 0.5 part |
| Vectashield | 140 | 1 part |
| Citifluor | 750 | 5.5 part |

### 3. Product information

| Product | Description | Company | Art.Nr.: | Size |
|---|---|---|---|---|
| **Alexa picolyl Azide** | Click-iT™ Plus Alexa Fluor™ 488 Picolyl Azide Toolkit | Thermo Fisher Scientific | C10641 | 1 kit |
| **Citifluor** | Glycerol/ PBS solution AF1 | Citifluor Ltd. (Electron microscopy science) | 17970 | 100 mL |
| **DAPI** | DAPI stain | Sigma-Aldrich | D9564 | 10 mg |
| **DMSO** | Dimethyl sulfoxide for molecular biology (DMSO) | Sigma-Aldrich | D8418-50ML | 50 mL |
| **Gelatin** | Gelatin from bovine skin Type B, powder, BioRegent, suitable for cell culture | Sigma-Aldrich | G9391 | 100G |
| **HPG** | Click-IT™ L-Homopropargylglycine (HPG) | Thermo Fisher Scientific | C10186 | 5 mg |
| **Polycarbonate filter** | 0.2 µm, 25 mm diameter | Millipore | GTTP02500 | 100 pcs |
| **Support filter** | 0.45 µm, 25 mm diameter | Millipore | HAWP02500 | 100 pcs |
| **Vectashield** | | Vector Laboratories, Inc. | H-1000 | 10 mL |
| **Water** | Sigma Water | Sigma-Aldrich | W4502 | 1 L |

**Additional lab equipment, materials:**

- Blotting paper, tissue paper
- Centrifuge conical tubes (50 mL)
- Ethanol for cleaning (70%)
- Fluorescence microscope (1000x magnification), Immersion oil
- Forceps, Scalpel
- Hybridization oven
- Microcentrifuge tubes (0.6 mL, 1.5 mL)
- Permanent marker, Pencil
- Petri dish
- Slide glass, Cover slip (24 x 60 mm)
- Water bath

### 4. Image analysis with ACMEtool3 (Bennke et al. 2016)

**ACMEtool3:**

https://www.mpi-bremen.de/en/automated-microscopy.html#section19794
https://ada-scientific.ch/technobiology/acmetool.html

**Procedures:**

*** Blue text indicates the typical image analysis settings used in our lab. These can be adjusted depending on image quality and the specifications of the microscope.**

1. Prepare TIF, 8-bit, grayscale microscopy images in a folder (usually, TIF is better quality than JPEG).
2. Choose '**1. Analyze Directory'**
3. Select the folder where the images are
4. Go to **B)** Check the photos to see if they are good.
5. Go to '**C) Image processing'**
6. Correlate '**Reference Image'** to '**Image Processing Method**'.
   Before pressing '**Add**', '**Corresponding parameters'** should be carefully checked and adjusted if necessary.
   e.g. **DAPI channel** default setting for our lab's microscopic image is:
   **'Dynamic threshold'**, **Kernel Size: 21, Offset: 11**. If small cells are not detected with the default setting, adjust the **Kernel size to 19 and the offset to 7-9 (always an odd number)**. Also, **uncheck** '**Remove regions**'. You can also change the '**Channel Name'** and **colour.** If you are OK with the setting, press '**Add**'.
   The next channel is BONCAT-HPG (referred to as the **FITC channel**); we usually use the same settings as for the DAPI setting. These units are in pixels.
7. Start image processing and save the file. (The information about corresponding parameters cannot be viewed after creating the metadata file. Therefore, if I change the Kernel size and Offset from the default, I will include this information in the filename.) Go to '**2. Load Metadata (*.IM3)'** to open the file
8. '**Menu**' '**Expand Tree**' to see all the images.
9. Go to the '**Info/Settings**' tab, and if needed, change the '**Counting Frame'.** We usually use the default **20-pixel** offsets.
10. Go back to '**FOV Browser**'—first **DAPI channel**. Check '**set**' to see the detection of the cells. Similarly, check the **FITC channel** as well. This is just to see the first detection, which usually needs to be adjusted (next step).
11. Go to '**Set Definitions**' and adjust the setting to suit your sample.
    For example, the default setting of our microscopic pictures is:
    **DAPI channel, Set Definition: Area>12 and Area<300 and SBR>1.2**
    **Sub Set Definition: Nr_FITC_Signals>0**
    **FITC channel, Set Definition: Area>6 and Area<300**
    **Subset Definition: Area>6**
    **From DAPI to FITC, Overlap Definition: Percent >0**
    These settings also depend on the resolution/quality of the images, which can vary across different labs. Thus, some manual inspections would help decide the setting.

Press '**Calculate valid cells**'. To view the detection results, go to '**FOV Browser**' to see the images. Repeat this process until you are satisfied with the settings.

12. If you want to know a specific single cell information, go to '**FOV Browser**' '**Show**' '**Cell Info**' and then choose '**Select**'. This information can be used to modify the '**Set Definition**', whether you wish to include or exclude the target cells.
13. If you are satisfied with your setting, review all the DAPI images. If you find a poor-quality image, uncheck it to exclude it from the analysis.
14. Similarly, review all the FITC images.
15. In the '**FOV Browser**', you can also check the '**DAPI**' -> '**subset**' to see whether you like the detection of HPG-positive DAPI.
16. After you are fine with the cell detection, go to '**Reports**'
17. To get cell counts per image, go to '**Features Sample/ FOV Report**'. The numbers displayed as output are indicated in **green text**. You can activate or deactivate by clicking the text.
18. Press '**Generate FOV Report**'.
19. If only a summary report is needed, press '**Generate Sample Report**'
20. Go to '**Menu**' -> **Save** or copy the output. Paste into a spreadsheet.

**Additional info ('Cell Info'):**

- Area: detected area in pixels
- Circularity: roundness (1 = perfectly round)
- Elongation: length/width
- SBR: signal to background ratio
- MGV = mean gray value (brightness of object, 0: Black, 255: White, in 8-bit images)
- MVGp90: mean of 10% brightest pixels
- MGVp10: mean of 10% darkest pixels
- MGV bg: mean gray value of the local surrounding

**For single-cell intensity analysis:**

- Perform '**1 Analyze Directory and Create Metadata**' with the images. Use identical settings if the images are already analyzed for either the Sample Report or the FOV Report (count data). For the intensity analysis, note that the **FITC** channels are placed as the first channel and **DAPI** as the second channel in the '**Channel List**'.
- After making the IM3 file, go to '**2. Load Metadata (*.IM3)**'. Apply the same settings as cell counts for '**Set Definition**', then press '**Calculate valid cells**'.
- Go to '**Report**' -> '**Features Cells Report**', choose **Sub Set** for '**Selection**' and **FITC** for '**Channel**', so that we can get FITC cell reports which are DAPI positive.
- Press '**Generate CELL Report**'.
- Save the output data.
